# Spectrum graph-based de-novo sequencing algorithm MaxNovo achieves high peptide identification rates in collisional dissociation MS/MS spectra

**DOI:** 10.1101/2021.09.04.458985

**Authors:** Petra Gutenbrunner, Pelagia Kyriakidou, Frido Welker, Jürgen Cox

## Abstract

We describe MaxNovo, a novel spectrum graph-based peptide de-novo sequencing algorithm integrated into the MaxQuant software. It identifies complete sequences of peptides as well as sequence tags that are incomplete at one or both of the peptide termini. MaxNovo searches for the highest-scoring path in a directed acyclic graph representing the MS/MS spectrum with peaks as nodes and edges as potential sequence constituents consisting of single amino acids or pairs. The raw score is a sum of node and edge weights, plus several reward scores, for instance, for complementary ions or protease compatibility. For search-engine identified peptides, it correlates well with the Andromeda search engine score. We use a particular score normalization and the score difference between the first and second-best solution to define a combined score that integrates all available information. To evaluate its performance, we use a human cell line dataset and take as ground truth all Andromeda-identified MS/MS spectra with an Andromeda score of at least 100. MaxNovo outperforms other software in particular in the high-sensitivity range of precision-coverage plots. We also identify incomplete sequence tags and study their statistical properties. Next, we apply MaxNovo to ion mobility-coupled time of flight data. Here we achieve excellent performance as well, except for potential swaps of the two amino acids closest to the C-terminus, which are not well resolved due to the low end of the mass range in MS/MS spectra in this dataset. We demonstrate the applicability of MaxNovo to palaeoproteomics samples with a Late Pleistocene hominin proteome dataset that was generated using three proteases. Interestingly, we did not use any machine learning in the construction of MaxNovo, but implemented expert domain knowledge directly in the definition of the score. Yet, it performs as good as or better than the leading deep learning-based algorithm.

## Introduction

De-novo sequencing^1,2^ has the goal of determining the amino acid sequence of a peptide directly from its tandem mass spectrum without making use of a peptide sequence database. Potential fields of applications in proteomics are widespread and include the study of ancient samples^3^, identification of HLA peptides^4–6^, monoclonal antibody sequencing^7,8^, and detection of endogenous, non-ribosomal peptides^9^. From early on it was found advantageous to represent the spectrum as a graph^2^ in which the fragment peaks correspond to nodes which are connected by edges, whenever the mass difference is interpretable as a mass of one or more amino acids, and hence the connected peaks are adjacent in an ion series. Optimal paths in these graphs are then determined with diverse computational methods including dynamic programming^10^, hidden Markov models^11^ and probabilistic network modeling^12^. More recently, deep learning was applied to the problem^13–17^, which has led to the best performing methods to date.

The MaxNovo algorithm described in this manuscript makes use of the spectrum graph representation as well. It performs an exhaustive search for the best path using a cost function that is designed in a way such that the resulting score is similar to the Andromeda^18^ search engine score, in cases where the database search led to an identification as well. MaxNovo is fully integrated in MaxQuant^19,20^ which allows to make use of results obtained from the search engine-based workflow, as for instance the accurate calibrated precursor masses and the three-dimensional MS1 features and isotope patterns. On an Orbitrap HeLa benchmark dataset, we show that MaxNovo performs well in terms of total number of correct de-novo identifications with controlled number of wrong identifications. It outperforms other software, as, for instance PEAKS^21^, in the high specificity region. Further studies on tims-TOF Pro data, as well as an ancient proteomics application, demonstrate MaxNovo’s applicability to diverse mass spectrometry proteomics data sets.

## Experimental Procedures

### HeLa dataset

Mass spectrometric raw data from a HeLa cell line was obtained from the PRIDE repository PXD006932^22^. What we refer to as ‘single-shot’ dataset are the three biological replicates measured on a Q Exactive HF-X using an Orbitrap resolution setting of 15000 (Thermo Fisher Scientific, Bremen, Germany). ‘Fractionated data’ are the 46 fractions obtained by Q Exactive HF-X using an Orbitrap resolution setting of 7500. The MaxQuant runs for both the single shot and the fractionated data were done against the *H. sapiens* proteome (UP000005640), including isoforms, and downloaded from UniProt on 07/04/2021. In total six separate MaxQuant runs were performed for the single shot data. One run with all five scores (complement score, protease score, water-loss score, ammonia-loss score, a_2_ score) contributing to the calculation of the raw score and other five runs with each one of the score contributions omitted. For the fractionated data only one MaxQuant run was made with the default calculation of the raw score including all five scores. Default values for all parameters were used in the MaxQuant analysis.

### Tims-TOF pro dataset

HeLa DDA data acquired on a tims-TOF Pro instrument was downloaded from the PRIDE repository PXD022582^23^ and was searched with MaxQuant against the *H. sapiens* proteome (UP000005640) including isoforms downloaded from UniProt on 07/04/2021. In the MaxQuant analysis for all parameters default values were used. To switch on tims-TOF analysis the parameter ‘Type’ was set to TIMS-DDA.

### Ancient dataset

Mass spectrometric raw data from a hominin bone specimen was obtained from the PRIDE repository PXD018264^24^. The three biological replicates from the samples digested with the proteases trypsin, chymotrypsin, and Glu-C, were grouped per protease and searched in MaxQuant in three separate runs using different proteome databases. One MaxQuant run was searched against the *H. sapiens* proteome (UP000005640) downloaded from UniProt on 05/08/2021. Two subsequent MaxQuant runs were made, one against the *Gorilla gorilla gorilla* proteome (UP000001519) and one against the *Pan troglodytes* (UP000002277), both downloaded from UniProt on 04/08/2021. All three Uniprot databases contain one protein sequence per gene. In the MaxQuant analysis the default settings were applied except for the following settings: No fixed modification was selected and as variable post-translational modification oxidation of Methionine, deamidation of asparagine and glutamine, hydroxylation of proline, and carbamidomethylation of cysteine. The minimum peptide length was set to eight (default value is seven) and the minimum score for unmodified peptides was set to 40 (default is 0). The Glu-C specificity was configured to also include C-terminal cleavage after glutamine (Q), alongside C-terminal cleavage of glutamic acid (E) and aspartic acid (D) as it was defined in the original publication.

### Pre-processing of MS/MS spectra for MaxNovo search

All MS/MS spectra are subject to the standard pre-processing that is also applied before submitting spectra to the Andromeda search. Isotope patterns are found based on correlation to the averagine model^25^. In case at least two peaks were put together in an isotope pattern, the corresponding monoisotopic peak replaces them with the summed intensity of the member peaks. In case the charge determined from the isotope pattern is larger than one, the peak is added as a singly charged version. If more than one charge state was found for a fragment (within a user definable tolerance), these are summed up into a single peak.

### NOVOR data preparation and analysis

The single-shot and fractionated HeLa raw files were converted to the mgf file format with ProteoWizard^26^ version 3.0.11579 for processing with Novor^27^ (Version v1.06.0634, Java SDK 16). The protease is trypsin, which is the only supported option. HCD was selected as fragmentation method and FT as mass analyzer. The precursor error tolerance was set to 15 ppm and the error tolerance for fragment ions to 0.02 Da.

### PEAKS data analysis

Both HeLa datasets, single-shot and fractionated, were analyzed by the de-novo algorithm in the PEAKS software^28^ (version PEAKS X Pro, Peaks Studio 10.6 build 20201221). For the de-novo search, the instrument ‘Orbitrap (Orbi-Orbi)’ was selected which has set by default the parent mass error tolerance to 15 ppm and the fragment mass error tolerance to 0.02 Da. Trypsin was defined as digestion enzyme and its cleavage specificity was configured that it cleaves after arginine and lysine also if a proline follows. As fixed modification carbamidomethylation of cysteine was included and as variable modifications oxidation of methionine and acetylation of protein N-term. For the maximal number of variable PTMs the default value of three is selected. We increased the number of reported candidates per spectrum from five up to ten. The parameter “Feature association for chimera scans” in the data-refinement section was deactivated, since co-eluting peptides are not within the scope of our benchmark setup. For the top scoring de-novo sequence comparison, the de-novo sequence with the highest de-novo score was selected. In case of multiple de-novo sequences having the same top de-novo score, all sequences are taken as top sequence.

### Benchmark based on the HeLa datasets

For both HeLa datasets, single-shot and fractionated, as ground truth the identified MS/MS spectra based on the Andromeda database search engine at 1% PSM and 1% Protein FDR (default settings) were taken which were further filtered by an Andromeda score greater than or equal to 100. All isoleucine in the database sequence as well as de-novo sequence were replaced by leucine. The de-novo identified MS/MS spectra of each of the tools MaxNovo, PEAKS and Novor are joined based on the raw file name and scan number information.

### BLAST search

The de-novo sequence identifications, which were not identified by the Andromeda database search engine, were validated by an iterative local BLAST (version 2.11) search. Only completely de-novo sequenced MS/MS spectra were considered and filtered to have a combined score of at least 91.715. Next, for each de-novo sequence all combinations, up to a maximal number of 300, were generated and submitted to four separate BLAST searches each using a different database. For the first three searches blastp was performed by setting the following parameters: window_size = 40, word_size = 2, evalue 1000, max_hsps = 1, - threshold 11 and against one of the following databases: 1) Swiss-Prot filtered by only human, 2) Swiss-Prot filtered by only *Bos Taurus* and 3) Swiss-Prot. The fourth search was a tblastn search against a manually generated HeLa nucleotide database based on RNA-seq data ^29,30^ (ERR127306_1.fastq, ERR127306_2.fastq, ERR127307_1.fastq, ERR127307_2.fastq). All BLAST results were filtered to have for each submitted de-novo sequence identification the best BLAST hit. In case a de-novo sequence identification was validated by multiple database hits the following database priority was applied: 1) Swiss-Prot human, 2) HeLa RNA-seq, 3) Swiss-Prot *Bos taurus* and 4) Swiss-Prot.

### Software and data availability

MaxNovo is integrated into the MaxQuant software from version 2.0.3.0 onward and can be downloaded from https://www.maxquant.org/maxquant/. A user guide on how to run MaxNovo in MaxQuant is provided as part of the Supporting Information, Supplementary Table 1 contains a list of MaxQuant parameters relating to MaxNovo with their explanations. Supplementary Table 2 describes all new MaxNovo associated columns in the ‘msmsScans.txt’ output file.

## Results and Discussion

### MaxNovo spectrum graph

Input for the MaxNovo algorithm are individual MS/MS spectra in which the fragment peaks are de-isotoped and transformed to charge state one (Fig. 1a), in the same way as MS/MS spectra are prepared in MaxQuant for the peptide database search with Andromeda ^18^ (see Experimental Procedures). Hence, the MaxNovo algorithm assumes that fragment ions are singly charged. The precursor masses are derived from three-dimensional features, spanned by m/z, retention time and signal intensity, after the standard nonlinear mass calibration in MaxQuant has been applied, leading to high accuracy mass estimates for the precursor ions. For each MS/MS spectrum, we construct a graph with the peaks in the spectrum as nodes (Fig. 1b). The nodes that correspond to peaks in the MS/MS spectrum we call ‘internal nodes’. Four additional nodes are added to the graph, which represent the N-terminus and the C-terminus each twice, once for being reached at low masses, corresponding to the beginning of an ion series, and once for being reached at high masses, corresponding to the end of an ion series.

**Fig. 1:**
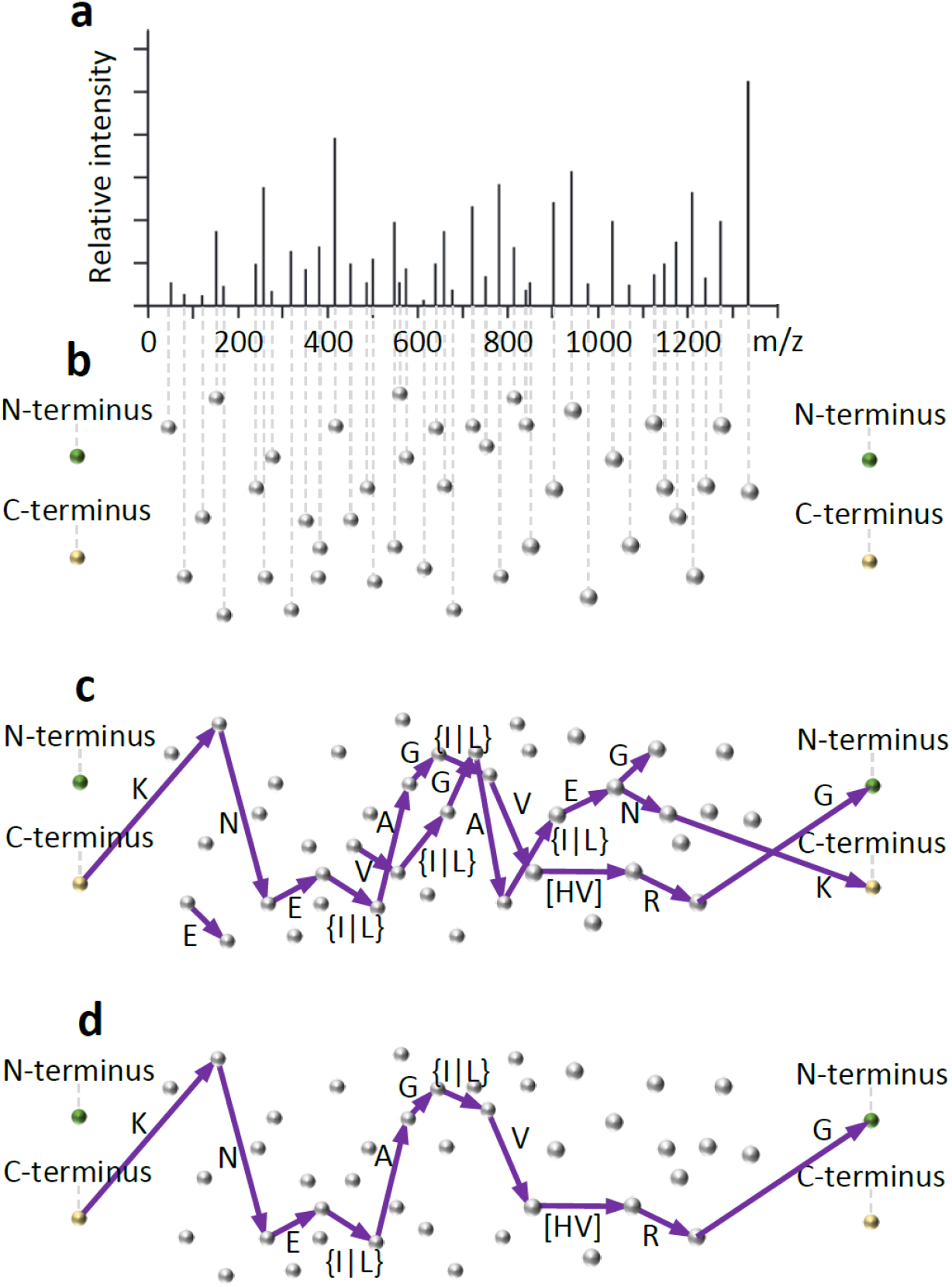
Schematics of the MaxNovo spectrum graph. **a**. Starting point for the algorithm is the processed MS/MS spectrum with de-isotoped fragments transferred to charge equals one, and the accurate precursor mass obtained from three-dimensional MS1 features after their nonlinear recalibration. **b**. Each peak in the processed spectrum becomes a potential ‘internal node’ of the spectrum graph. Four ‘terminal nodes’ are added, corresponding to N- or C-terminus reached at low or high masses. **c**. A directed edge from low to high mass is put between two nodes, whenever their mass difference fits single or paired amino acids. **d**. The path with the highest raw score is found in an exhaustive search. It may start and end with terminal or external nodes. The raw score is the sum of weights of traversed nodes and edges plus several reward scores as described in the main text.

We then build a directed acyclic graph (DAG) by placing edges between vertices from lower to higher mass. This is based on the 20 common amino acids, and modified versions of some amino acids, which can be specified by the user as fixed and variable modifications. An edge is placed between two internal nodes if their mass difference fits a single amino acid mass or the sum of two amino acid masses (Fig. 1c). Mass steps that have equal mass or are so close in mass that they are not discernible based on the data are grouped together and treated as the same amino acid (as, for instance leucine and isoleucine). Similarly, as for internal nodes, for the termini we connect the beginning node with internal nodes that can be reached with a single or a double amino acid step, considering the appropriate terminal masses for the y or b ion series. It is only known to which ion series (e.g. y or b) a path through the network belongs by arriving at one of the terminal nodes. By default, we require a mass difference between two edges to match with a maximal error of 25 ppm. Then weights are assigned to edges and nodes as defined in the next subsection. In this graph, we perform an exhaustive search for the path with the highest raw score (Fig. 1d) based on a recursive algorithm. Either the best path represents the y or the b series, depending on which of the two achieves a better score. In case the path with the highest raw score does not reach the termini at one end, a second search for a best path is performed, with the constraint to fill the missing mass. This is necessary for the case that y and b series are not overlapping.

### MaxNovo raw score

Each path through the spectrum DAG gets a raw score assigned. The path in this graph with the highest raw score is defined to be the result of the de-novo search for this MS/MS spectrum and this highest score is the MaxNovo raw score for this spectrum. Either the best path can connect both termini, in which case the complete peptide has been sequenced, or it can be incomplete at one or both of the termini. Hence, the MaxNovo algorithm is a combination of complete de-novo sequencing algorithm and sequence tag finding algorithm within one unified scoring scheme. The raw score assigned to each path consists of six contributions

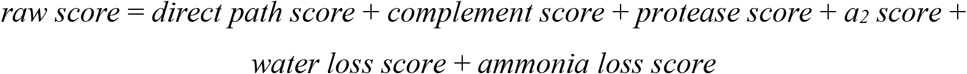

which correspond to specific rewards or penalties. The *direct path score* is the sum of scores defined on the edges and nodes that constitute the path and is scoring one main ion series, which is supposedly the one that contributes most to the identification. Therefore, the path represents either part of, or the complete b-series or part of, or the complete y-series. It is not supposed to mix contributions from N-terminal or C-terminal series, since it receives contributions from a path consisting of steps that correspond to single amino acid or amino acid pair mass differences within one ion series. Two-amino-acid steps are allowed in order to be able to have complete solutions connecting the termini also when one or several peaks in a series are missing. By default, we allow up to two two-steps in a path. Each node visited by the path contributes -log_10_ ((*g* + 1) / 100) to the score, where *g* is the number of peaks in a 100 Da interval centered on the current peak which have a higher intensity than the current peak. This contribution assigns a higher reward to peaks with a high intensity relative to other peaks in the surrounding 100 Da interval. A traversed edge contributes -log_10_ (*s*), where *s* is the number of potential steps of the type that was actually taken that could have been taken from the current node. This can have two values: if a single step was taken, then it is the total number of single steps available, and if a two-step was taken, this is the total number of two-steps. Essentially, this is a penalty on how many mass differences were tried in order to reach the next node. For instance, there are more potential two-steps than there are single-steps, which get down-weighted accordingly. Overall, a path length dependent score contribution of -log_10_(*n*) is added to the total direct path score, where *n* is the total number of steps in the path.

The *complement score* looks for the presence of nodes corresponding to complementary ions to the ions found in the direct path. For instance, if the path describes b-ions, these complementary ions correspond to y-ions that each match with one of the b-ions as a complementary pair. The roles of y and b ions could also be the opposite. For each complementary peak found, a contribution -log_10_ ((*g* + 1) / 100) / 4 is added, similarly as the node weight in the direct path score. Additionally, if a complementary ion is found that resolves the order of the amino acids in a two-step of the direct path a corresponding score contribution is added that equals to the situation as if the resolving peak was found in the direct path. The *protease score* adds a reward in case the path reaches a terminus at which an expectation is met regarding the protease used for digesting proteins to peptides in the sample preparation. For instance, in the case of trypsin, a path reaching the C-terminus would be rewarded if the last step were either an arginine or a lysine, in case it is a single amino acid step, or it contains these amino acids, in case it is a two amino acid step. The *a*_*2*_ *score* adds a contribution in case the a_2_ ion is found. This is only possible in case the sequence path is continued to the N-terminus in order to know that it follows the b-series. The *water loss score* checks for the presence of one of the amino acids D, E, S or T in the path traversed so far. In case any of these is present, it is checked if peaks are present at the expected position(s) for water losses. The *ammonia loss score* similarly checks for peaks at the characteristic masses for ammonia losses based on the presence of K, N, Q or R in the so far identified sequence.

For convenience, we provide brief definitions of these scores and scores defined in subsequent subsections in Table 1. In Fig. 2 we show a scatter plot of the raw score and the Andromeda score for identified spectra in the single-shot HeLa dataset, which have a Pearson correlation of 0.71, which reduces to 0.60 when no Andromeda score filter is applied. In the construction of the raw score, the main purpose was that it reflects well the total evidence that is present in a spectrum for the peptide sequence that it claims to have found. This is confirmed by the good correlation with the Andromeda score.

**Table 1:** Definition of scores used in this publication. The scores with their names in italic are additive contributions to the raw score. The combined score is the score of choice for ranking full-length sequences. Incomplete sequence tags are best ranked using the normalized score.

|  |  |
| --- | --- |
| raw score | This is the score assigned to each path, which is the sum of the six following score. The optimal path is found maximizing this score in an exhaustive search over all paths. |
| *direct path score* | Score summing up weights that are collected along the path with contributions from traversed nodes and edges. |
| *complement score* | Score contribution of ions that are complementary to the ions in the direct path. |
| *protease score* | Reward for terminal parts of the sequence that are in agreement with the specified protease that was used for generating peptides from proteins. |
| *a2 score* | Reward for the presence of an a2 ion. |
| *water loss score* | Reward for the presence of ions resulting from the loss of a water molecule in case the main path contains any of the amino acids D, E, S or T. |
| *ammonia loss score* | Reward for the presence of ions resulting from the loss of an ammonia molecule in case the main path contains any of the amino acids K, N, Q or R. |
| normalized score | This is the raw score divided by the precursor mass. While the raw score adds up the total spectral evidence for a peptide, the normalized score rather corresponds to the spectral evidence per peptide length. |
| complete score | The complete score equals the normalized score, in case the sequence goes from terminus to terminus, i.e. is completely sequencing the peptide. Otherwise the complete score equals zero. |
| gap score | The gap score is the difference in raw scores between the best and the second-best scoring solution. If there is no second-best solution, the gap score equals the raw score. |
| combined score | The combined score is a combination of the complete score and the gap score. Both of these scores are ranked, Then the sum of the two ranks is taken and normalized to lie between 0 and 100. |

**Fig. 2:**
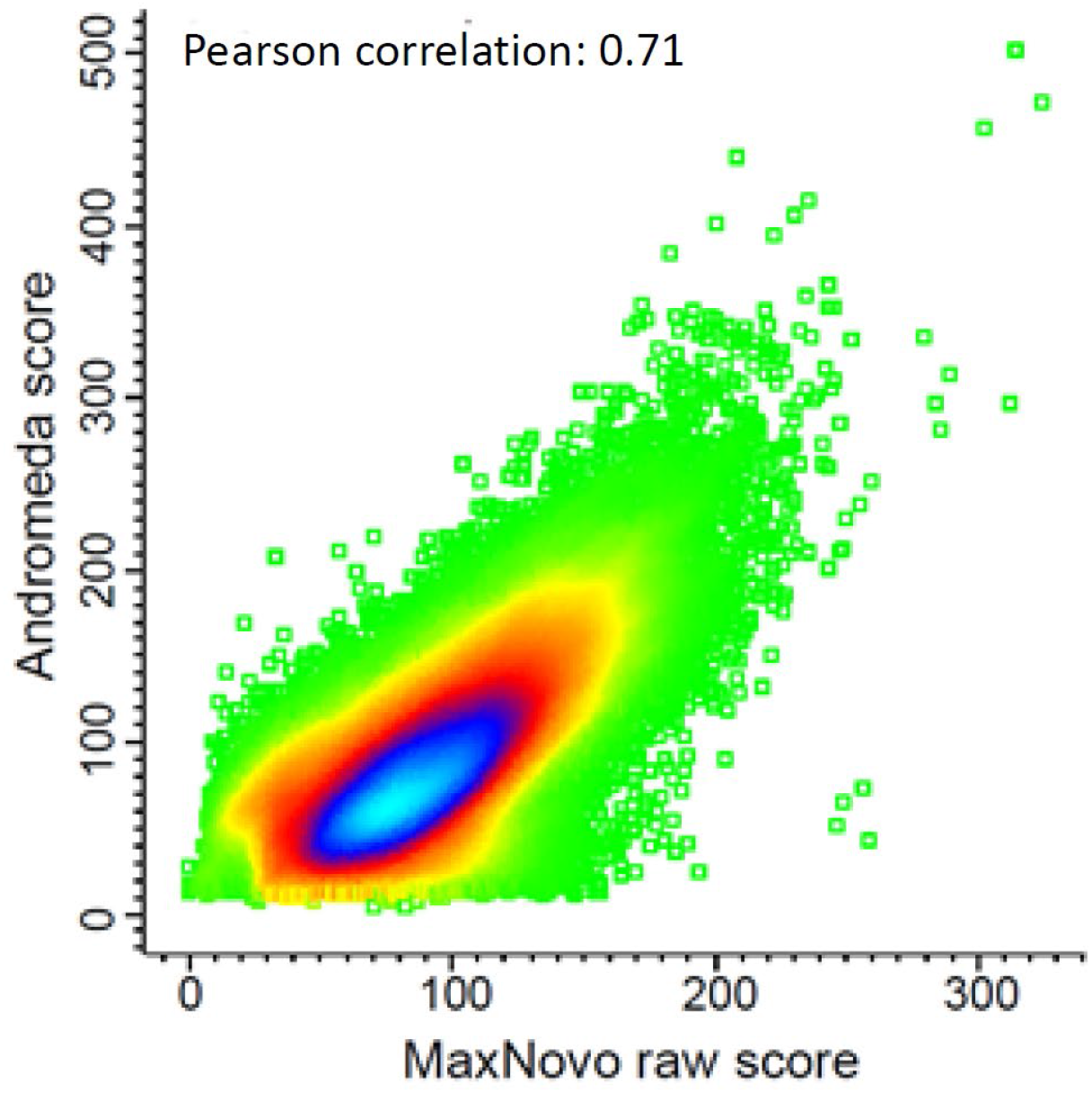
Scatter plot of the raw score against the Andromeda score. Data is filtered for Andromeda score > 100. Data points are colored by their density. The Pearson correlation is 0.71.

### Normalized score, gap score and combined score

The raw score introduced in the previous section is optimized for finding the best solution, either a sequence tag or a full-length sequence, for a given MS/MS spectrum. However, it is not made for comparing two optimal solutions in two different MS/MS spectra. The ability to compare scores across different spectra is, however, crucial for obtaining confidence in an identification and for estimating the percentage of false positives. To see this in a better way, we define as a dataset with essentially known ground truth all MS/MS spectra in the HeLa dataset that were identified in the MaxQuant analysis with default protein and PSM level false discovery rates (FDRs) of 1% each, and additionally filter the spectra to have an Andromeda score of at least 100. We consider these spectra as our ground truth and the aim of the de-novo sequencing algorithm is to identify as many peptide sequences of these correctly, if possible the complete sequence for each, and otherwise as many amino acids as possible in a sequence tag. For now, we restrict ourselves to identifying complete peptide sequences.

We will frequently use precision-coverage plots, which are created by ranking the spectra according to a given score. A de-novo identification is counted as correct only if it completely agrees with the Andromeda sequence. If there is only one deviation, the whole peptide counts as incorrect. For a given score threshold we define precision as the number of correctly identified sequences divided by all spectra with a score above the threshold. With coverage, we mean the number of correctly identified spectra above the threshold divided by the total number of spectra in the ground truth. Our definition of coverage measures how many spectra from the whole ground truth have been correctly identified with a complete sequence.

If we calculate such a precision-coverage plot for the raw score defined in the previous subsection (green curve in Fig. 3a) we see that the performance is not ideal. In particular, in the high specificity range on the left side, no good precision values are achieved. In order to fix this problem, we define the normalized score, which is the raw score divided by the precursor mass. It is a measure for sequence evidence per length. In particular, in a situation where there is good sequence evidence only in parts of the sequence and lack of it otherwise, the normalized score will not be particularly high. Indeed, the normalized score has better precision-coverage characteristics (purple curve in Fig. 3a). As ‘complete score’, we define it to be equal to the normalized score in case it suggests a full-length peptide sequence and zero otherwise (blue curve in Fig. 3a). Another aspect of potential relevance for judging the correctness of a de-novo sequence is how well the second-best solution scores compared to the best. If the score difference between the two highest solutions is small, there is a certain likelihood that the second-best solution is the correct one, while if this gap is large, this strengthens the plausibility of the top-scoring solution. The precision-coverage curve based on this gap score (red curve in Fig. 3a) results in a similar area under the curve as the complete score. Since the gap score and the complete score measure different aspects of the identification, it should be beneficial to combine the two scores. Indeed, we define the combined score as the sum of the two ranks of the complete score and the gap score and it achieves the best precision-coverage characteristics of all scores (orange curve in Fig. 3a). The combined score is hence the method of choice for finding complete sequences, and it is implicitly meant when referring to the MaxNovo score without further specification.

**Fig. 3:**
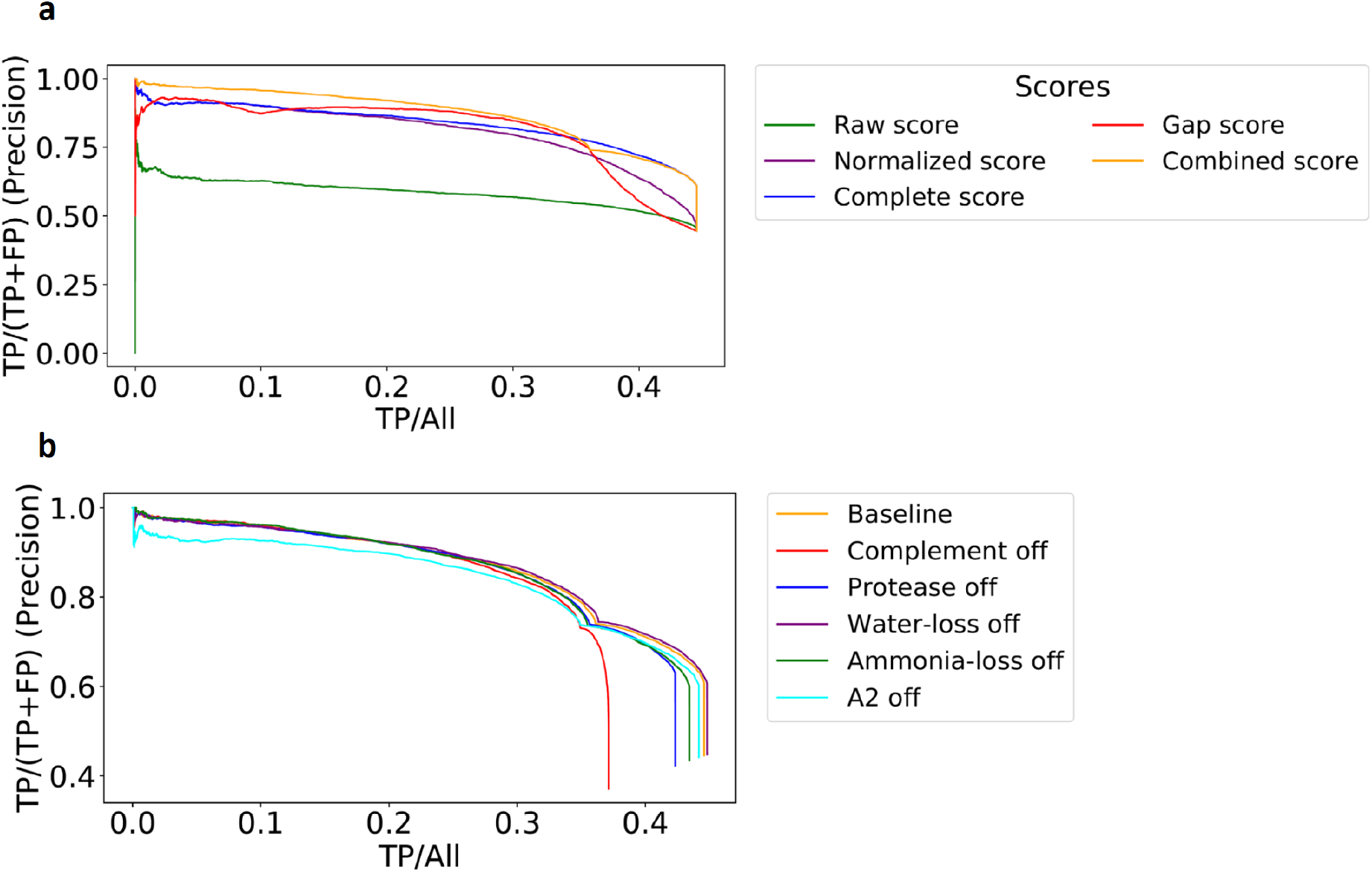
Precision-coverage plots of several MaxNovo scores. **a**. Data was obtained from the single-shot HeLa dataset with all identified MS/MS spectra with an Andromeda score of at least 100. Only full peptide sequences without any amino acid mistake are counted as correct. The curves correspond to raw score (green), normalized score (purple), complete score (blue), gap score (red) and combined score (yellow). **b**. The combined score is shown again as baseline (color). For the other curves, each time one score contribution has been omitted.

Next, we investigated what the benefits are of the individual additive contributions to the raw score is to the overall performance. As the baseline, we take the precision-coverage curve based on the combined score (yellow curve in Fig. 3b). Then we remove each of the contributions to the raw score except the direct path score one at a time and record precision-coverage curves for these as well. In the high-specificity region, the strongest effect came from the scoring of the a_2_ ion. Conversely, the inclusion of the complementary ion series (the b-series ions in case the main series scored by the direct-path score is the y series) made a big difference in the low specificity region. Water loss, ammonia loss and protease score all contribute in less significant and similar amounts.

### Benchmark and comparison to other software

In order to compare the performance of MaxNovo to other software we analyzed the same data with the PEAKS and Novor algorithms (see Experimental Procedures). Fig. 4 shows precision-coverage plots for all three programs. While Novor is less performant, the other two are close in performance. Up to coverage 0.35, the curve for MaxNovo is on top. The PEAKS curve reaches farther to the right end, meaning it finds slightly more sequences with low certainty and the respective areas under the curves are very similar for PEAKS and MaxNovo. We conclude that these two software platforms are showing comparable performances with MaxNovo performance being better in the higher specificity region.

**Fig. 4:**
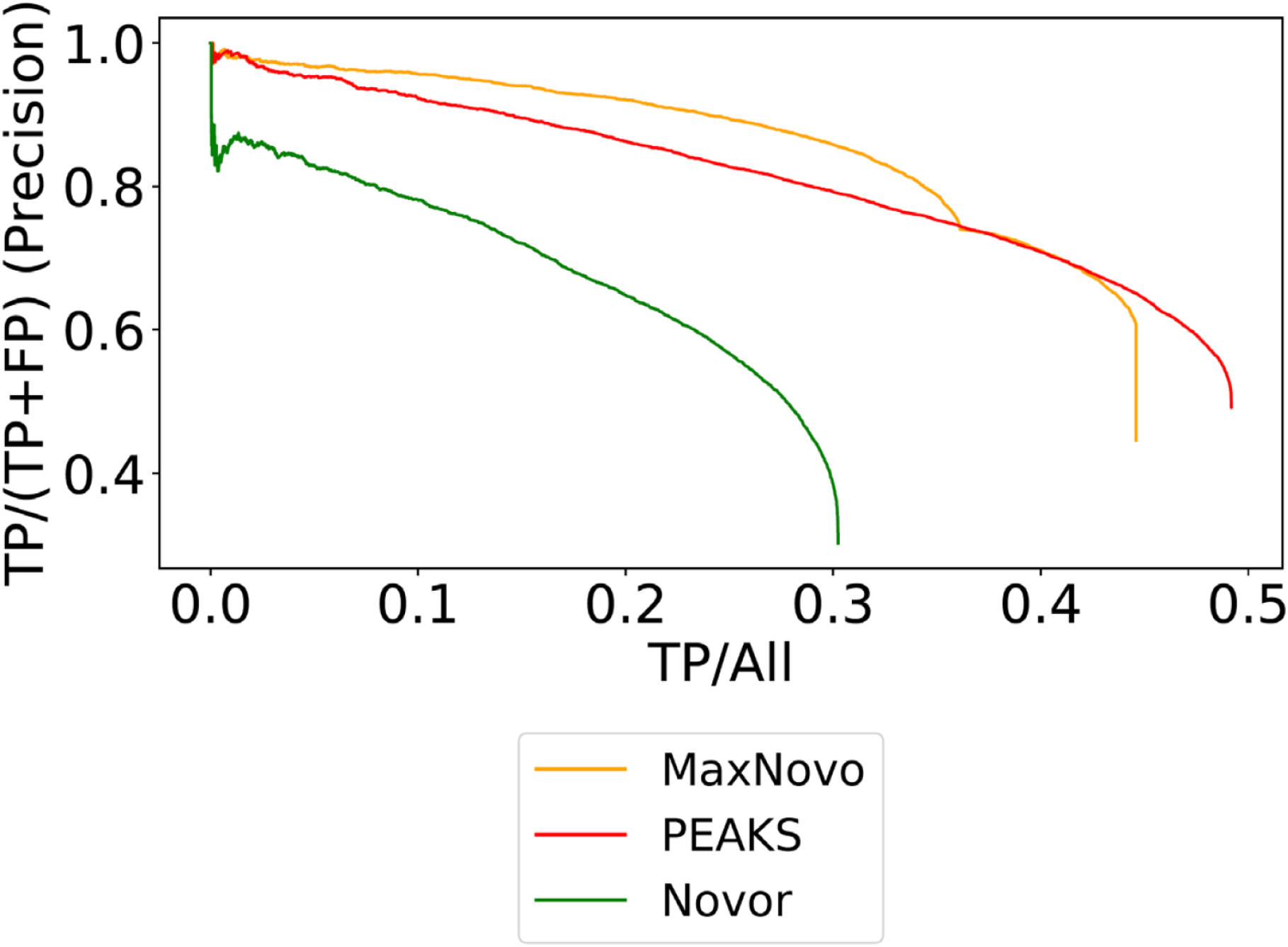
Performance comparison of the combined score. Precision-coverage curve comparing the combined score of MaxNovo with PEAKS and Novor.

### Incomplete sequences

To measure the performance on incomplete sequences, we recorded precision-coverage plots using the normalized score and we counted as correct, when the predicted sequence agrees completely with the Andromeda sequence or a sub-sequence of it (Fig. 5a). This we do for the single shot and the fractionated HeLa datasets separately. We find that we obtain curves that are similar to each other above a precision of 0.8 and are deviating below that. Next, we determine what the expected precision of the results is at a given normalized score threshold (Fig. 5b). The curves for fractionated and single-shot data agree well in particular at larger score values, which indicates that the dependency of the precision on the score could be generalizable. For instance, a precision of 0.8 corresponds to a normalized score threshold of approximately ten.

**Fig. 5:**
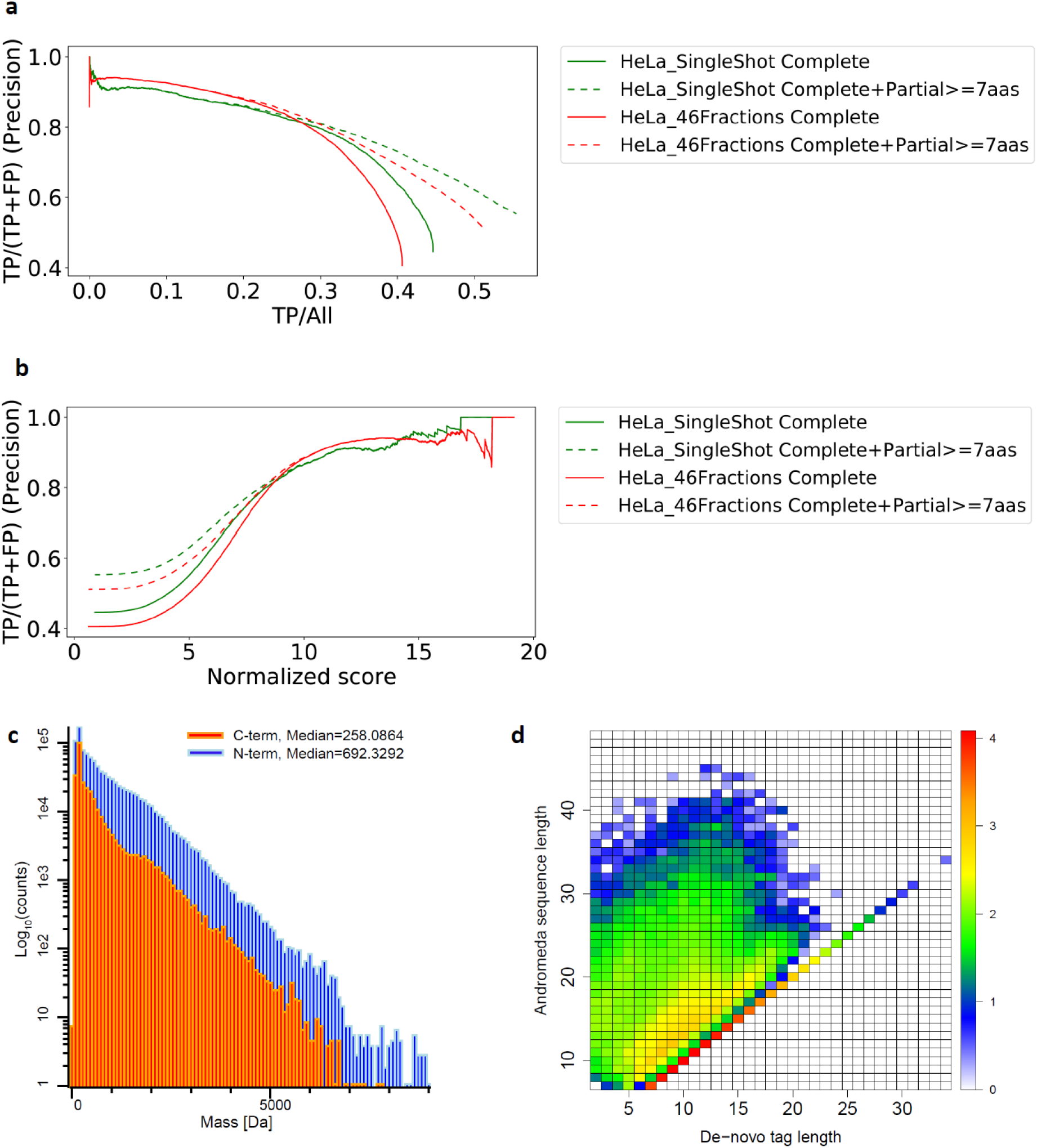
Incomplete sequences. **a**. Precision-coverage curve based on the MaxNovo normalized score including complete and incomplete sequences calculated for the single-shot and the fractionated dataset. A minimum tag length of seven was applied. **b**. The precision values from panel a are plotted against the normalized score. **c**. Histograms of N- and C-terminal missing masses for the fractionated HeLa dataset (all MS/MS spectra with missing terminal mass > 0). **d**. Density plot of tag length vs. peptide sequence length of all MS/MS spectra that have also been identified by Andromeda.

Fig. 5c shows histograms of N- and C-terminal missing masses, having medians of 692.3 and 258.1 Da, respectively in the fractionated dataset. The y-axis of the histogram is in logarithmic scale and one can see that higher masses are much less frequent. A density scatter plot of peptide sequence length vs. tag length (Fig. 5d) is dominated by the values on the diagonal corresponding to full-length sequences. The distribution of partial sequences has a median length of twelve.

### New identifications

Next, we focus on the sequences in the fractionated HeLa dataset that have not been identified by the Andromeda search engine. 15.2%, or 135,908 in total, of the complete sequences or sequence tags that are at least seven amino acids long are from MS/MS spectra that have not been identified by Andromeda. We perform a multi-tier BLAST search against protein databases containing all human proteins, proteins derived from the HeLa genome, the *Bos taurus* proteome and finally against a Swiss-Prot database containing all species. We further filtered the non-identified MS/MS spectra by a combined score of 91.715 to allow an error rate of 10%, which results in 28,126 sequences that are submitted to BLAST. 28,123 sequences find a match with the same length in at least one of the four tiers. Out of these, 18,492 (65.8%) are matching exactly, while the remaining ones have at least one substitution (Fig. 6a). We further classify the sequences without mismatch in Fig. 6b.

**Fig. 6:**
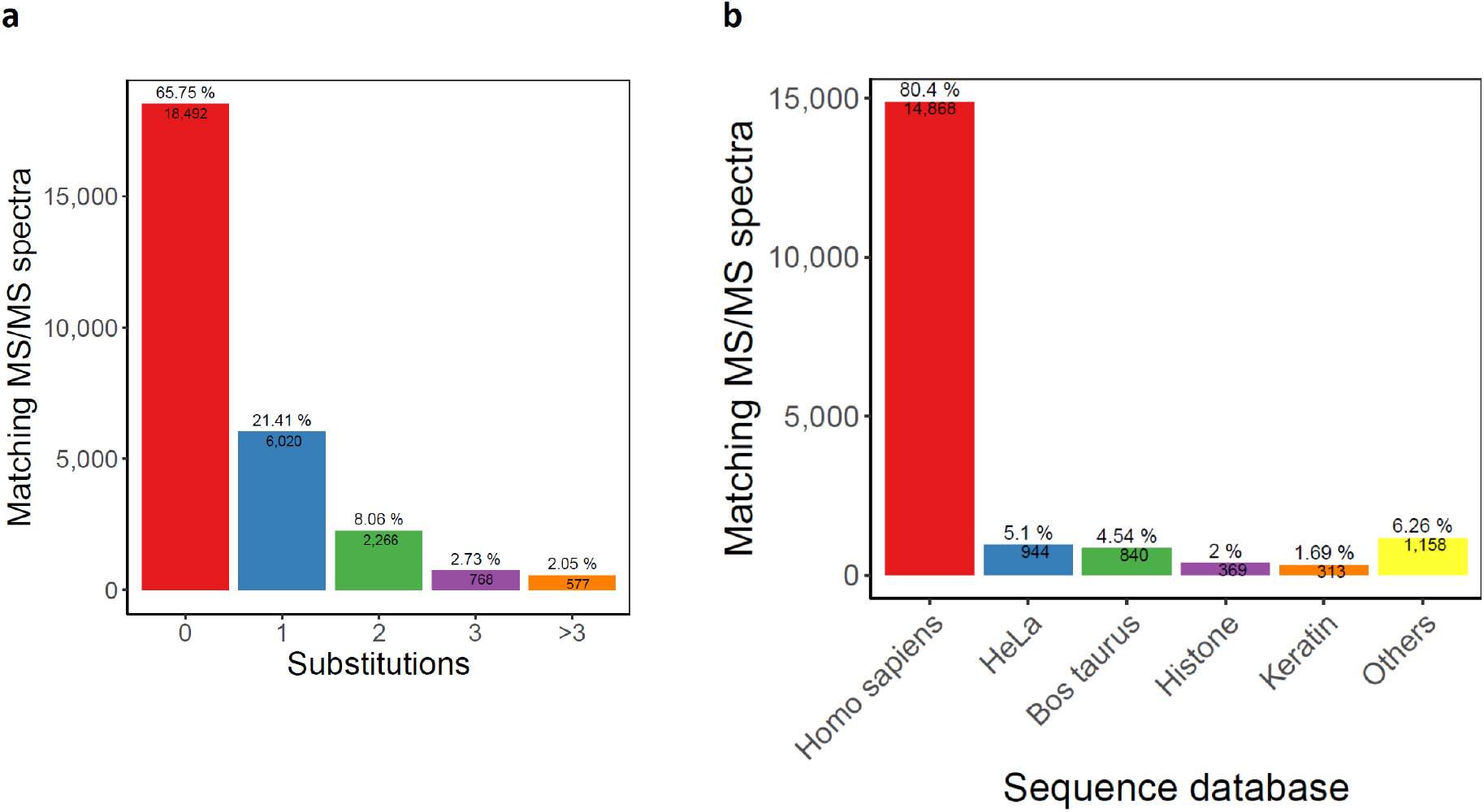
New identifications. **a**. Number of MS/MS spectra in the fractionated HeLa dataset whose full-length de-novo prediction was matched by BLAST with the specified number of substitutions. **b**. The de-novo sequences from panel a that were matched without substitutions are classified according to the sequence database that they were matching to.

### Tims TOF data

We next applied MaxNovo to a HeLa dataset acquired on a tims-TOF Pro instrument. The precision-coverage plot based on the combined score (Fig. 7) shows at first less performance than in Fig. 3. However, most of the wrong de-novo sequences have only a swap of the two C-terminal amino acids and are otherwise correct. This is due to the lower mass range limit in the MS/MS spectra of 200 Da in this particular dataset. If we ignore the order of the two C-terminal amino acids, we obtain a much better precision-coverage curve (Fig. 7). Hence, it is recommended to use a lower mass range limit for MS/MS spectra or to ignore the order of the last two amino acids in each peptide.

**Fig. 7:**
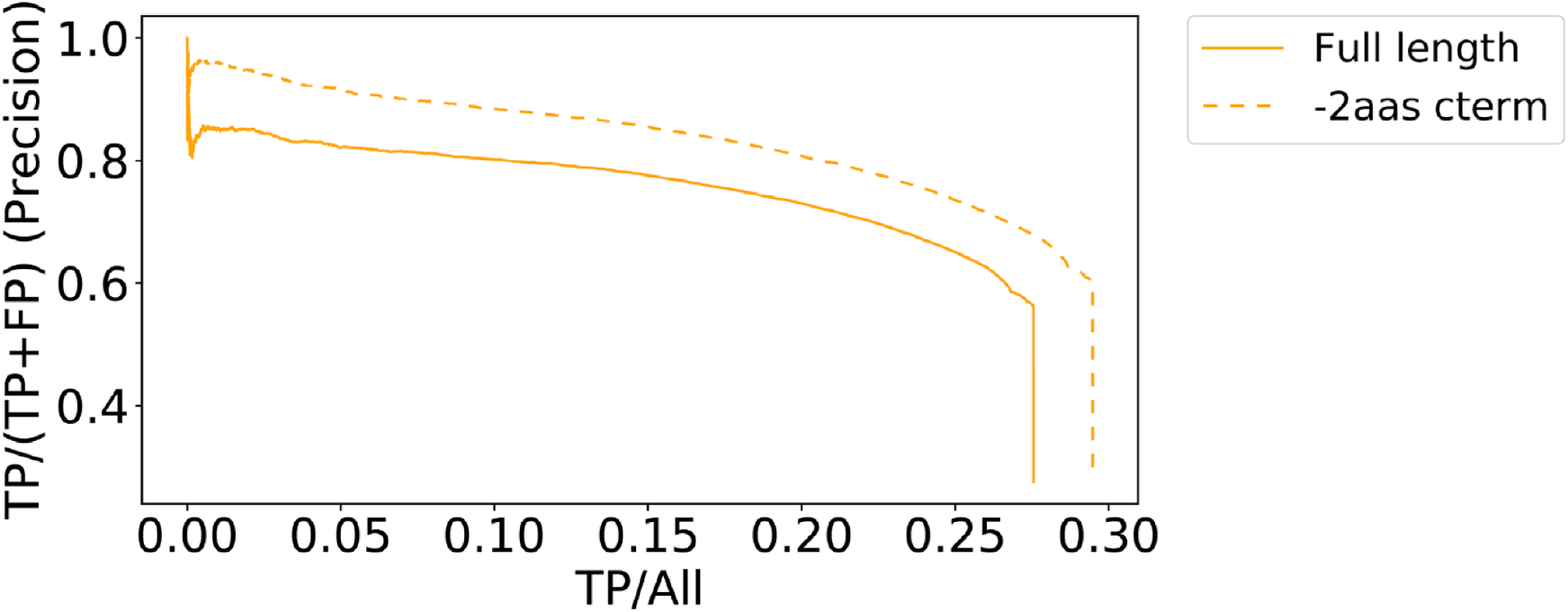
timsTOF Pro data. Precision-coverage curve for the timsTOF Pro data based on the combined score once checking complete correctness of the sequence and once ignoring the two C-terminal amino acids.

### Application to ancient sequences

Ancient proteomes present a challenging application of de-novo and error-tolerant search approaches, both because comparative genomes of closely-related species are frequently not available and because the determination of sequence variation is essential when phylogenetic analysis is the main purpose of the proteomic study. Ancient proteomes contain a large number of variable PTMs^31^, many of which might only be present at low frequencies, and have increasing rates of peptide bond hydrolysis for older samples, making the occurrence of semi-specific and non-specific peptides in resulting datasets much more likely^32^. Finally, dentine and bone proteomes are dominated by collagen type I, which is heavily hydroxylated, increasing search complexity as well as the presence of incomplete fragmentation series, particularly around proline positions. We therefore tested MaxNovo performance against a Late Pleistocene hominin proteome dataset that was previously generated using three proteases in parallel^24^. We observe that, as for modern data (Fig. 5d), de-novo sequence tag lengths are on average shorter when compared to the Andromeda-derived sequence solution (Fig. 8a). Likewise, as with PEAKS^33^, the probability that a top-ranking full-length MaxNovo solution exists and is correct for a given spectrum is highly dependent on peptide amino acid length, although the MaxNovo scores provide the possibility to apply stringent selection criteria and partly remove this dependency (Fig. 8b). Despite this, MaxNovo correctly resolved the true peptide sequence when presented with protein sequence databases of chimpanzee (*Pan*, Fig. 8c) and gorilla (*Gorilla*, Fig. 8d), including PSMs containing diagenetic modifications (deamidation) and proline hydroxylation.

**Fig. 8:**
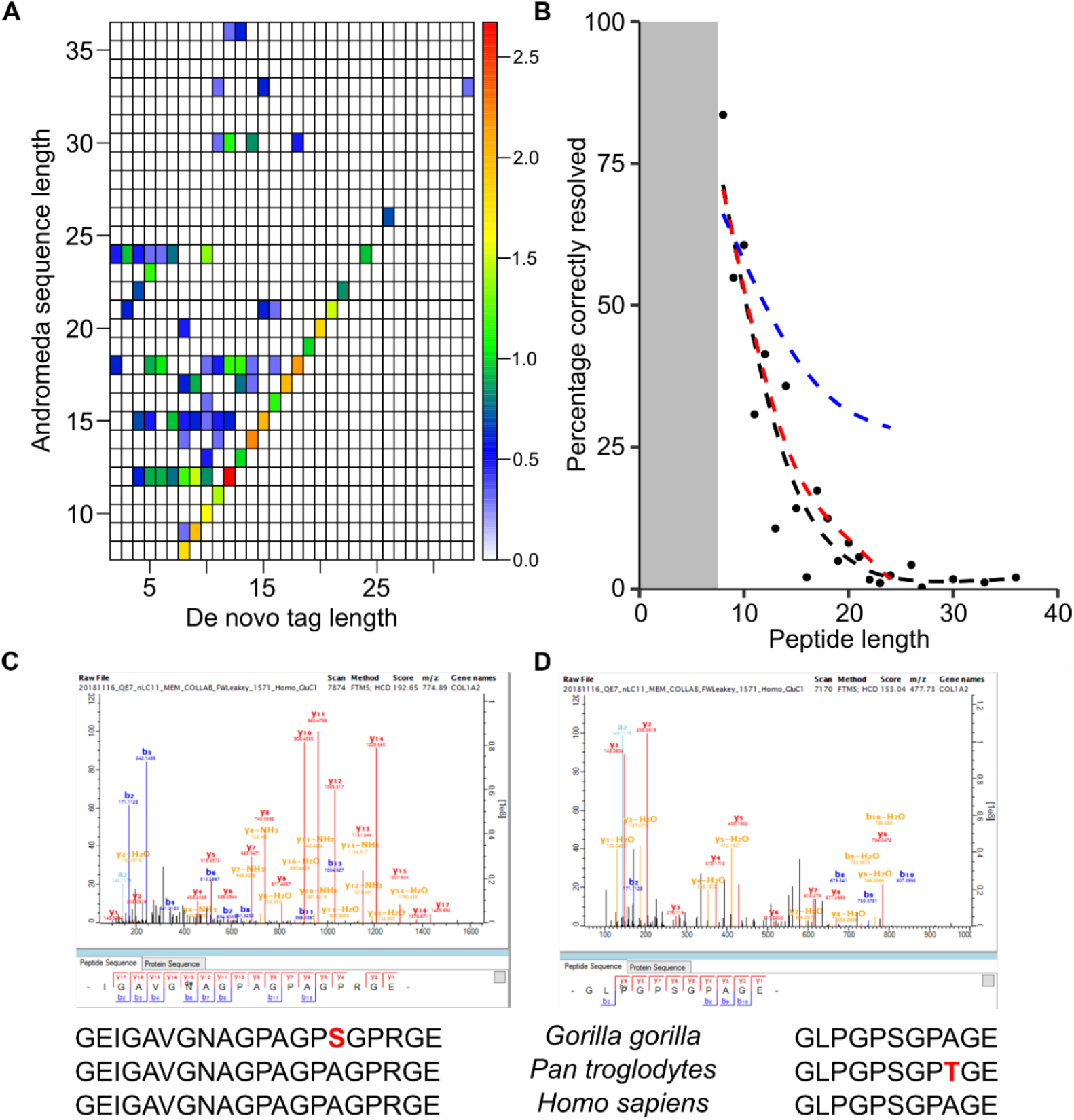
Validation on ancient samples. **a**. Density plot of tag length (MaxNovo) vs. peptide sequence length (Andromeda) for PSMs where the de-novo solution contains the database search sequence solution (n=1,726). **b**. Correctness of MaxNovo top-ranking sequence solutions are highly dependent on peptide sequence length (black, n=14,249). Filtering for combined scores over 75 increases average correctness moderately for longer peptides (red, n=5,703), but significantly when only taking into account normalized scores > 10 (blue, n=1,427). **c**. Example of a successfully resolved MaxNovo PSM containing asparagine deamidation (combined score = 98.9451, GluC digest). **d**. Example of a successfully resolved MaxNovo PSM containing proline hydroxylation (combined score = 98.907, GluC digest). For both c and d, a protein sequence alignment across COL1A2 is given, indicating relevant positions with SAP in *Gorilla* (c, A279S) and *Pan* (d, A564T). Coordinates in reference to UniProt accession number P08123.

## Conclusions

We introduced MaxNovo, a novel de-novo sequencing algorithm that is integrated into the MaxQaunt software. It shows a performance, in terms of sequence identifications, which is as good as or better than software that is currently in use. It is interesting to observe, that no machine learning was used in its construction, but that we rationalized a scoring function based on expert knowledge on peptide fragmentation spectra. Based on its integration into the MaxQuant environment, we can expect MaxNovo to make significant introductions to the recovery of peptide sequences of relevance in clinical, biological, and evolutionary settings.

## Acknowledgements

We thank Christoph Wichmann, Hamid Hamzeiy and Sule Yilmaz for helpful discussions. This project was partially funded by the German Ministry for Science and Education (BMBF) funding action MSCoreSys, reference number FKZ 031L0214D. P.K. is supported by the Marie Skłodowska-Curie European Training Network (ETN) PUSHH, a project funded by the European Union’s Framework Program for Research and Innovation Horizon 2020 (grant agreement no. 861389). P.G. is supported by the Marie Skłodowska-Curie European Training Network TEMPERA, a project funded by the European Union’s EU Framework Program for Research and Innovation Horizon 2020 under grant agreement no. 722606. F.W. is supported by the European Research Council (ERC) under the European Union’s Horizon 2020 research and innovation programme (grant agreement no. 948365).

## Author Contributions

P.G. and J.C. designed and developed the main code. F.W. provided and analyzed the hominin bone data. All authors analyzed the data and wrote the manuscript.

## References

(1) Taylor, J. A.; Johnson, R. S. Sequence Database Searches via de Novo Peptide Sequencing by Tandem Mass Spectrometry. Rapid Commun. Mass Spectrom. 1997, 11 (9), 1067–1075. https://doi.org/10.1002/(SICI)1097-0231(19970615)11:9<1067::AID-RCM953>3.0.CO;2-L.

(2) Dančík, V.; Addona, T. A.; Clauser, K. R.; Vath, J. E.; Pevzner, P. A. De Novo Peptide Sequencing via Tandem Mass Spectrometry. In Journal of Computational Biology; 1999. https://doi.org/10.1089/106652799318300.

(3) Cappellini, E.; Prohaska, A.; Racimo, F.; Welker, F.; Pedersen, M. W.; Allentoft, M. E.; De Barros Damgaard, P.; Gutenbrunner, P.; Dunne, J.; Hammann, S.; Roffet-Salque, M.; Ilardo, M.; Moreno-Mayar, J. V.; Wang, Y.; Sikora, M.; Vinner, L.; Cox, J.; Evershed, R. P.; Willerslev, E. Ancient Biomolecules and Evolutionary Inference. Annual Review of Biochemistry. 2018. https://doi.org/10.1146/annurev-biochem-062917-012002.

(4) Ternette, N.; Yang, H.; Partridge, T.; Llano, A.; Cedeño, S.; Fischer, R.; Charles, P. D.; Dudek, N. L.; Mothe, B.; Crespo, M.; Fischer, W. M.; Korber, B. T. M.; Nielsen, M.; Borrow, P.; Purcell, A. W.; Brander, C.; Dorrell, L.; Kessler, B. M.; Hanke, T. Defining the HLA Class I-Associated Viral Antigen Repertoire from HIV-1-Infected Human Cells. Eur. J. Immunol. 2016. https://doi.org/10.1002/eji.201545890.

(5) Khodadoust, M. S.; Olsson, N.; Wagar, L. E.; Haabeth, O. A. W.; Chen, B.; Swaminathan, K.; Rawson, K.; Liu, C. L.; Steiner, D.; Lund, P.; Rao, S.; Zhang, L.; Marceau, C.; Stehr, H.; Newman, A. M.; Czerwinski, D. K.; Carlton, V. E. H.; Moorhead, M.; Faham, M.; Kohrt, H. E.; Carette, J.; Green, M. R.; Davis, M. M.; Levy, R.; Elias, J. E.; Alizadeh, A. A. Antigen Presentation Profiling Reveals Recognition of Lymphoma Immunoglobulin Neoantigens. Nature 2017. https://doi.org/10.1038/nature21433.

(6) Laumont, C. M.; Vincent, K.; Hesnard, L.; Audemard, É.; Bonneil, É.; Laverdure, J. P.; Gendron, P.; Courcelles, M.; Hardy, M. P.; Côté, C.; Durette, C.; St-Pierre, C.; Benhammadi, M.; Lanoix, J.; Vobecky, S.; Haddad, E.; Lemieux, S.; Thibault, P.; Perreault, C. Noncoding Regions Are the Main Source of Targetable Tumor-Specific Antigens. Sci. Transl. Med. 2018. https://doi.org/10.1126/scitranslmed.aau5516.

(7) Bandeira, N.; Pham, V.; Pevzner, P.; Arnott, D.; Lill, J. R. Automated de Novo Protein Sequencing of Monoclonal Antibodies. Nature Biotechnology. 2008. https://doi.org/10.1038/nbt1208-1336.

(8) Tran, N. H.; Rahman, M. Z.; He, L.; Xin, L.; Shan, B.; Li, M. Complete de Novo Assembly of Monoclonal Antibody Sequences. Sci. Rep. 2016. https://doi.org/10.1038/srep31730.

(9) Bandeira, N.; Ng, J.; Meluzzi, D.; Linington, R. G.; Dorrestein, P.; Pevzner, P. A. De Novo Sequencing of Nonribosomal Peptides. In Lecture Notes in Computer Science (including subseries Lecture Notes in Artificial Intelligence and Lecture Notes in Bioinformatics); 2008. https://doi.org/10.1007/978-3-540-78839-3_16.

(10) Chen, T.; Tepel, M.; Rush, J.; Church, G. M.; Kao, M. Y. A Dynamic Programming Approach to de Novo Peptide Sequencing via Tandem Mass Spectrometry. J. Comput. Biol. 2001. https://doi.org/10.1089/10665270152530872.

(11) Fischer, B.; Roth, V.; Roos, F.; Grossmann, J.; Baginsky, S.; Widmayer, P.; Gruissem, W.; Buhmann, J. M. NovoHMM: A Hidden Markov Model for de Novo Peptide Sequencing. Anal. Chem. 2005. https://doi.org/10.1021/ac0508853.

(12) Frank, A.; Pevzner, P. PepNovo: De Novo Peptide Sequencing via Probabilistic Network Modeling. Anal. Chem. 2005. https://doi.org/10.1021/ac048788h.

(13) Tran, N. H.; Zhang, X.; Xin, L.; Shan, B.; Li, M. De Novo Peptide Sequencing by Deep Learning. Proc. Natl. Acad. Sci. 2017, 114 (31), 8247–8252. https://doi.org/10.1073/pnas.1705691114.

(14) Karunratanakul, K.; Tang, H. Y.; Speicher, D. W.; Chuangsuwanich, E.; Sriswasdi, S. Uncovering Thousands of New Peptides with Sequence-Mask-Search Hybrid de Novo Peptide Sequencing Framework. Mol. Cell. Proteomics 2019. https://doi.org/10.1074/mcp.TIR119.001656.

(15) Tran, N. H.; Qiao, R.; Xin, L.; Chen, X.; Liu, C.; Zhang, X.; Shan, B.; Ghodsi, A.; Li, M. Deep Learning Enables de Novo Peptide Sequencing from Data-Independent-Acquisition Mass Spectrometry. Nat. Methods 2019. https://doi.org/10.1038/s41592-018-0260-3.

(16) Yang, H.; Chi, H.; Zeng, W. F.; Zhou, W. J.; He, S. M. PNovo 3: Precise de Novo Peptide Sequencing Using a Learning-to-Rank Framework. In Bioinformatics; 2019. https://doi.org/10.1093/bioinformatics/btz366.

(17) Qiao, R.; Tran, N. H.; Xin, L.; Chen, X.; Li, M.; Shan, B.; Ghodsi, A. Computationally Instrument-Resolution-Independent de Novo Peptide Sequencing for High-Resolution Devices. Nat. Mach. Intell. 2021. https://doi.org/10.1038/s42256-021-00304-3.

(18) Cox, J.; Neuhauser, N.; Michalski, A.; Scheltema, R. A.; Olsen, J. V.; Mann, M. Andromeda: A Peptide Search Engine Integrated into the MaxQuant Environment. J. Proteome Res. 2011, 10 (4), 1794–1805. https://doi.org/10.1021/pr101065j.

(19) Cox, J.; Mann, M. MaxQuant Enables High Peptide Identification Rates, Individualized p.p.b.-Range Mass Accuracies and Proteome-Wide Protein Quantification. Nat Biotechnol 2008, 26 (12), 1367–1372. https://doi.org/10.1038/nbt.1511.

(20) Sinitcyn, P.; Tiwary, S.; Rudolph, J.; Gutenbrunner, P.; Wichmann, C.; Yilmaz, Ş.; Hamzeiy, H.; Salinas, F.; Cox, J. MaxQuant Goes Linux. Nat. Methods 2018, 15 (6), 401. https://doi.org/10.1038/s41592-018-0018-y.

(21) Ma, B.; Zhang, K.; Hendrie, C.; Liang, C.; Li, M.; Doherty-Kirby, A.; Lajoie, G. PEAKS: Powerful Software for Peptide de Novo Sequencing by Tandem Mass Spectrometry. Rapid Commun. Mass Spectrom. 2003, 17 (20), 2337–2342. https://doi.org/10.1002/rcm.1196.

(22) Kelstrup, C. D.; Bekker-Jensen, D. B.; Arrey, T. N.; Hogrebe, A.; Harder, A.; Olsen, J. V. Performance Evaluation of the Q Exactive HF-X for Shotgun Proteomics. J. Proteome Res. 2018, 17 (1), 727–738. https://doi.org/10.1021/acs.jproteome.7b00602.

(23) Sinitcyn, P.; Hamzeiy, H.; Salinas Soto, F.; Itzhak, D.; McCarthy, F.; Wichmann, C.; Steger, M.; Ohmayer, U.; Distler, U.; Kaspar-Schoenefeld, S.; Prianichnikov, N.; Yilmaz, Ş.; Rudolph, J. D.; Tenzer, S.; Perez-Riverol, Y.; Nagaraj, N.; Humphrey, S. J.; Cox, J. MaxDIA Enables Library-Based and Library-Free Data-Independent Acquisition Proteomics. Nat. Biotechnol. 2021. https://doi.org/10.1038/s41587-021-00968-7.

(24) Lanigan, L. T.; Mackie, M.; Feine, S.; Hublin, J. J.; Schmitz, R. W.; Wilcke, A.; Collins, M. J.; Cappellini, E.; Olsen, J. V.; Taurozzi, A. J.; Welker, F. Multi-Protease Analysis of Pleistocene Bone Proteomes. J. Proteomics 2020. https://doi.org/10.1016/j.jprot.2020.103889.

(25) Senko, M. W.; Beu, S. C.; McLaffertycor, F. W. Determination of Monoisotopic Masses and Ion Populations for Large Biomolecules from Resolved Isotopic Distributions. J Am Soc Mass Spectrom 1995, 6 (4), 229–233. https://doi.org/10.1016/1044-0305(95)00017-8.

(26) Chambers, M. C.; Maclean, B.; Burke, R.; Amodei, D.; Ruderman, D. L.; Neumann, S.; Gatto, L.; Fischer, B.; Pratt, B.; Egertson, J.; Hoff, K.; Kessner, D.; Tasman, N.; Shulman, N.; Frewen, B.; Baker, T. A.; Brusniak, M. Y.; Paulse, C.; Creasy, D.; Flashner, L.; Kani, K.; Moulding, C.; Seymour, S. L.; Nuwaysir, L. M.; Lefebvre, B.; Kuhlmann, F.; Roark, J.; Rainer, P.; Detlev, S.; Hemenway, T.; Huhmer, A.; Langridge, J.; Connolly, B.; Chadick, T.; Holly, K.; Eckels, J.; Deutsch, E. W.; Moritz, R. L.; Katz, J. E.; Agus, D. B.; MacCoss, M.; Tabb, D. L.; Mallick, P. A Cross-Platform Toolkit for Mass Spectrometry and Proteomics. Nat Biotechnol 2012, 30 (10), 918–920. https://doi.org/10.1038/nbt.2377.

(27) Ma, B. Novor: Real-Time Peptide de Novo Sequencing Software. J. Am. Soc. Mass Spectrom. 2015. https://doi.org/10.1007/s13361-015-1204-0.

(28) Zhang, J.; Xin, L.; Shan, B.; Chen, W.; Xie, M.; Yuen, D.; Zhang, W.; Zhang, Z.; Lajoie, G. A.; Ma, B. PEAKS DB: De Novo Sequencing Assisted Database Search for Sensitive and Accurate Peptide Identification. Mol. Cell. Proteomics 2012. https://doi.org/10.1074/mcp.M111.010587.

(29) Zarnack, K.; König, J.; Tajnik, M.; Martincorena, I.; Eustermann, S.; Stévant, I.; Reyes, A.; Anders, S.; Luscombe, N. M.; Ule, J. Direct Competition between HnRNP C and U2AF65 Protects the Transcriptome from the Exonization of Alu Elements. Cell 2013. https://doi.org/10.1016/j.cell.2012.12.023.

(30) Seo, J.; Singh, N. N.; Ottesen, E. W.; Lee, B. M.; Singh, R. N. A Novel Human-Specific Splice Isoform Alters the Critical C-Terminus of Survival Motor Neuron Protein. Sci. Rep. 2016. https://doi.org/10.1038/srep30778.

(31) Cappellini, E.; Welker, F.; Pandolfi, L.; Ramos-Madrigal, J.; Samodova, D.; Rüther, P. L.; Fotakis, A. K.; Lyon, D.; Moreno-Mayar, J. V.; Bukhsianidze, M.; Rakownikow Jersie-Christensen, R.; Mackie, M.; Ginolhac, A.; Ferring, R.; Tappen, M.; Palkopoulou, E.; Dickinson, M. R.; Stafford, T. W.; Chan, Y. L.; Götherström, A.; Nathan, S. K. S. S.; Heintzman, P. D.; Kapp, J. D.; Kirillova, I.; Moodley, Y.; Agusti, J.; Kahlke, R. D.; Kiladze, G.; Martínez-Navarro, B.; Liu, S.; Sandoval Velasco, M.; Sinding, M. H. S.; Kelstrup, C. D.; Allentoft, M. E.; Orlando, L.; Penkman, K.; Shapiro, B.; Rook, L.; Dalén, L.; Gilbert, M. T. P.; Olsen, J. V.; Lordkipanidze, D.; Willerslev, E. Early Pleistocene Enamel Proteome from Dmanisi Resolves Stephanorhinus Phylogeny. Nature 2019. https://doi.org/10.1038/s41586-019-1555-y.

(32) Chen, F.; Welker, F.; Shen, C. C.; Bailey, S. E.; Bergmann, I.; Davis, S.; Xia, H.; Wang, H.; Fischer, R.; Freidline, S. E.; Yu, T. L.; Skinner, M. M.; Stelzer, S.; Dong, G.; Fu, Q.; Dong, G.; Wang, J.; Zhang, D.; Hublin, J. J. A Late Middle Pleistocene Denisovan Mandible from the Tibetan Plateau. Nature 2019. https://doi.org/10.1038/s41586-019-1139-x.

(33) Welker, F. Elucidation of Cross-Species Proteomic Effects in Human and Hominin Bone Proteome Identification through a Bioinformatics Experiment. BMC Evol. Biol. 2018. https://doi.org/10.1186/s12862-018-1141-1.

