## Supporting Information for "Spectrum graph-based de-novo sequencing algorithm MaxNovo achieves high peptide identification rates in collisional dissociation MS/MS spectra"

### A user guide on how to run MaxNovo in MaxQuant

MaxNovo is integrated into the MaxQuant environment. You can download MaxQuant from https://maxquant.org/maxquant/ .

You must make sure to install .NET Core 3.1  SDK x64 from https://dotnet.microsoft.com/download/dotnet/3.1.

##### 1. Load your raw files and set your experiment design.

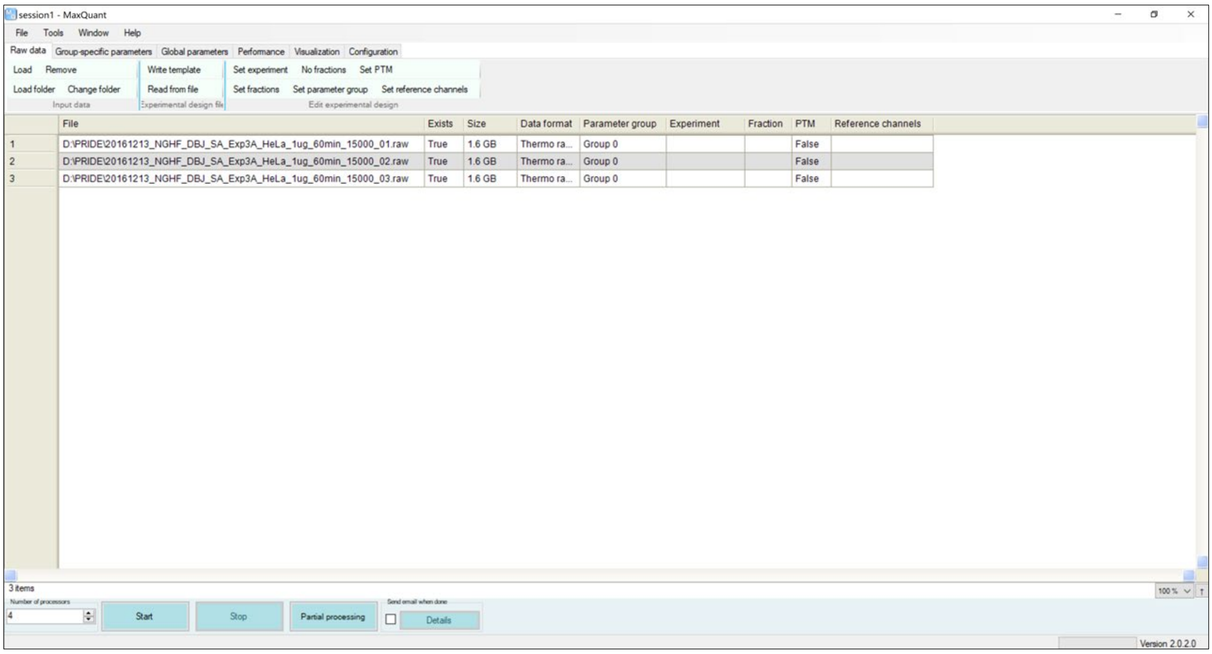

**2. Go to “Global parameters” tab → “Identification” tab.**

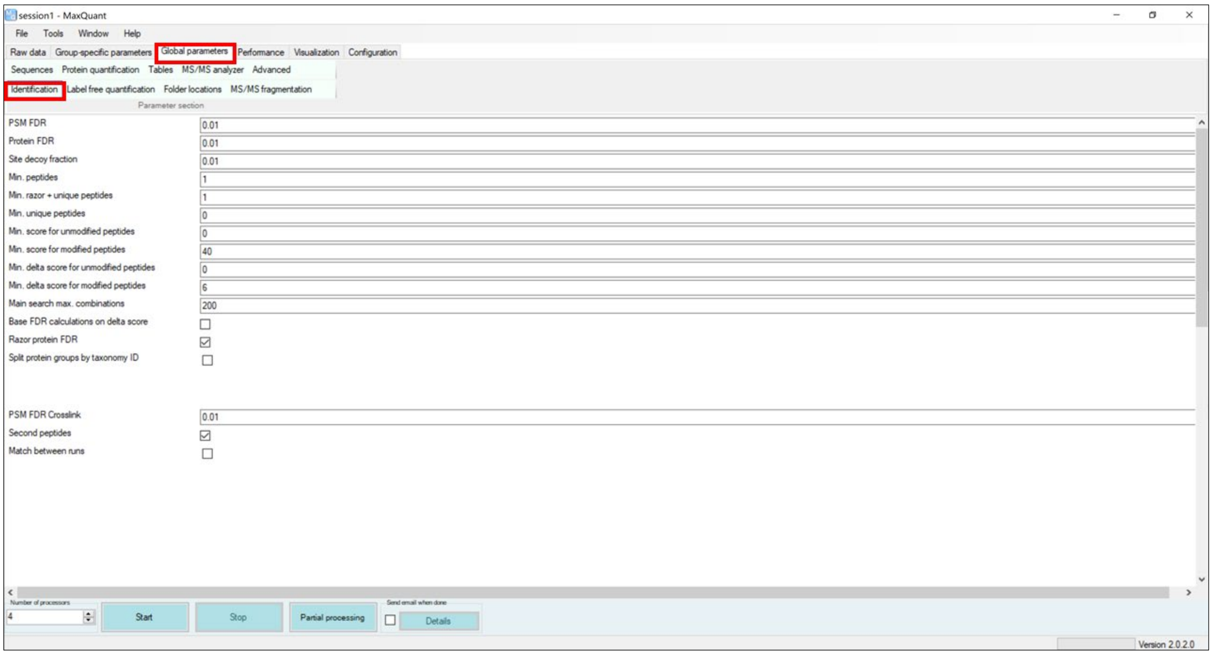

**2. a. Scroll down till you find the “De-novo sequencing” checkbox.**

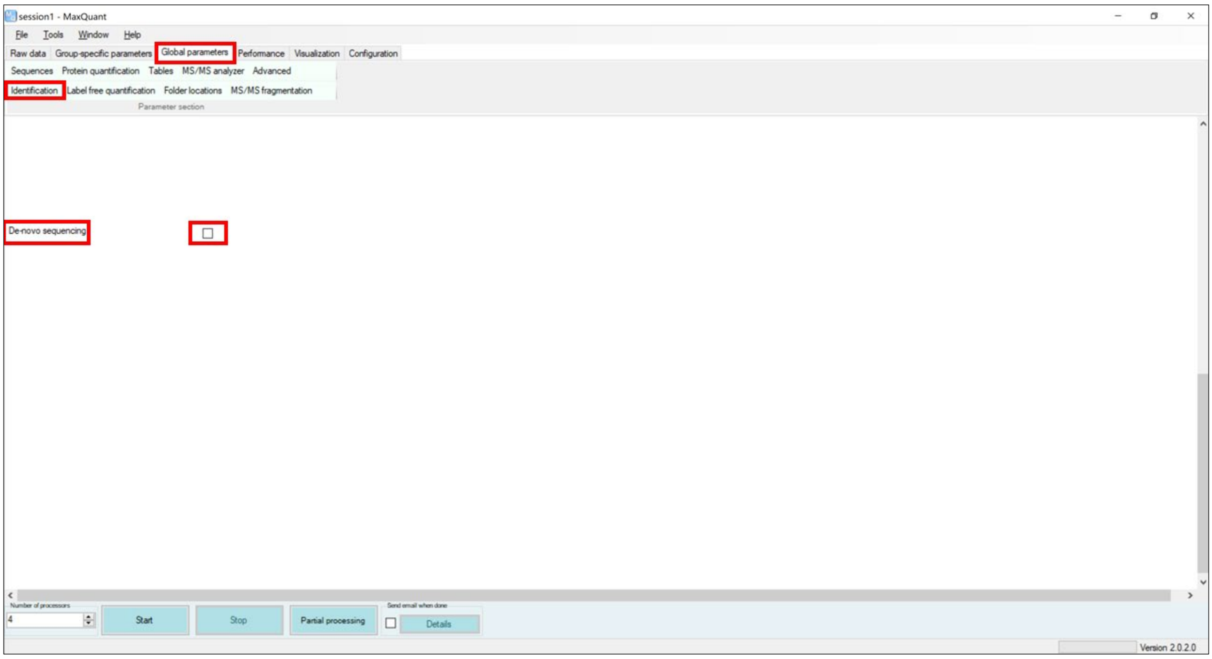

**2. b. Enable the De-novo sequencing by clicking on it.**

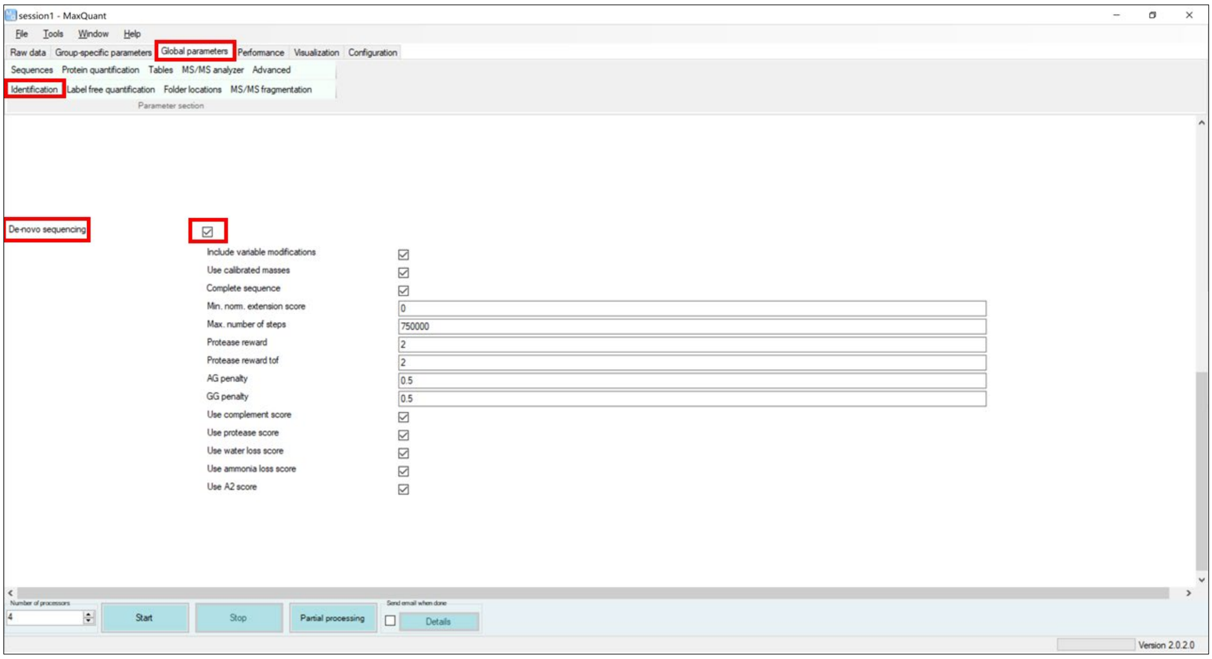

**2.c. Set the MaxNovo parameters that appeared after clicking on the "De-novo sequencing" checkbox.**

Please find the description of each one of the parameters at the Supplementary Table 1 in the Supporting Information:

**3. Press start to start a MaxQuant run with the MaxNovo denovo identification enabled.**

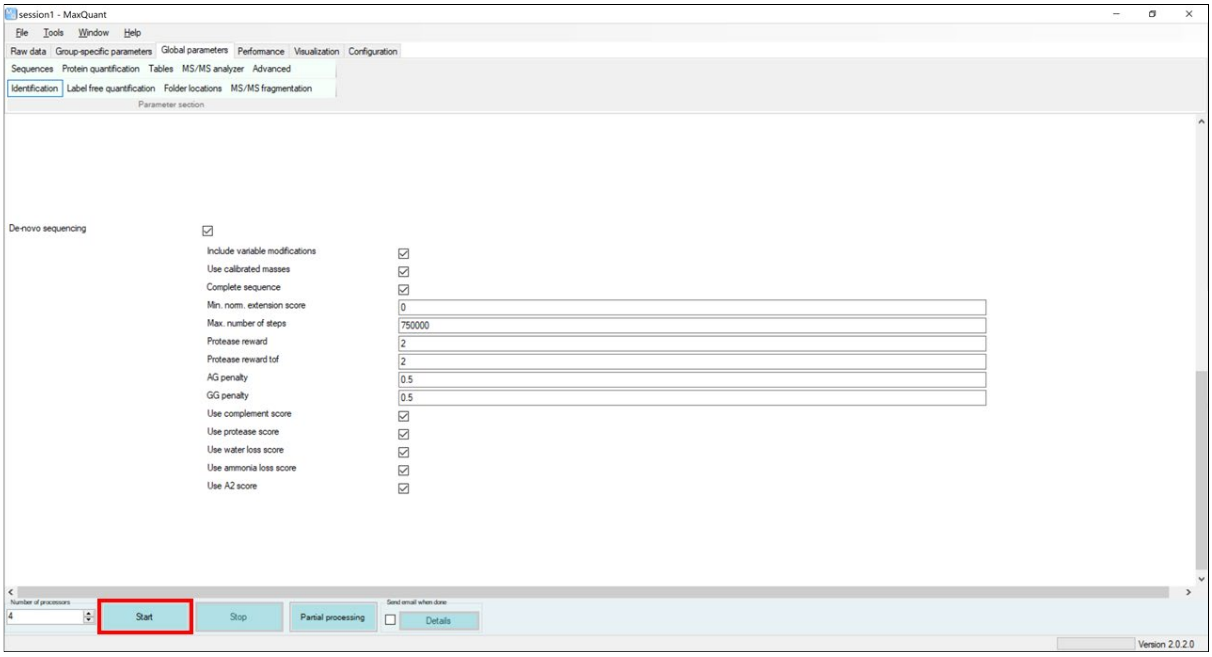

After MaxQuant finishes navigate to the “combined” folder that MaxQuant created and then to the “txt” folder. In the “txt” folder there is a file called “msmsScans.txt” where you can find all the additional de-novo information from your experiment. Please find the description of each one of the columns at the Supplementary Table 2 in the Supporting Information.

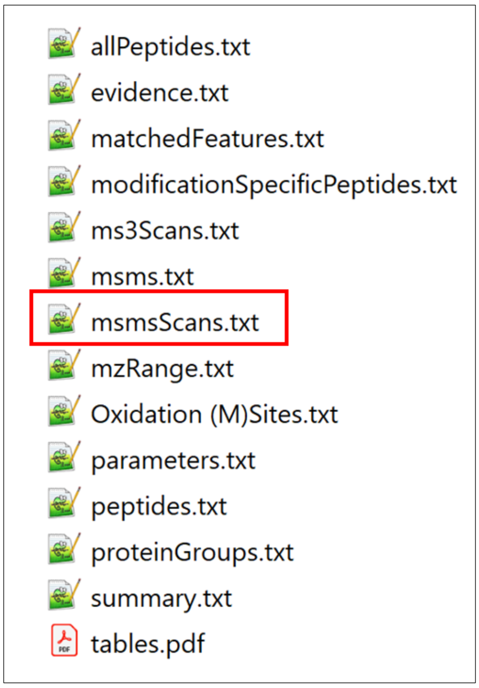

**Supplementary Table 1.**

Description of the de-novo parameters in MaxQuant. Find these parameters in MaxQuant when you enable the de-novo algorithm under Global parameters → Identification → De-novo sequencing.

| **Parameter** | **Default value** | **Description** |
|---|---|---|
| Include variable modifications | ✓ | When checked, the variable modifications (specified under Group-specific parameters -> Modifications -> Variable modifications) are considered when MaxNovo runs. |
| Use calibrated masses | ✓ | When checked, the masses are the ones after the re-calibration step. |
| Complete sequence | ✓ | When checked, if the de-novo sequence is not completely resolved, a complementary ion series is looked up that can extend it. |
| Min. norm. Extension score | 0 | the minimum normalized score the extended sequence has to have in order for the de-novo sequence to be extended (in case incomplete). |
| Max. number of steps | 750000 | The upper limit of steps in a path that are tested in the exhaustive search along the directed acyclic graph representing each MS/MS spectrum. |
| Protease reward | 2 | The weight of the contribution of the "protease score" in the calculation of the "raw score". |
| Protease reward tof | 2 | The weight of the contribution of the "protease score" in the calculation of the "raw score" for timsTOF data. |
| AG penalty | 0.5 | Sore penalty for picking AG instead of Q. |
| GG penalty | 0.5 | Sore penalty for picking GG instead of N. |

| Parameter | Default value | Description |
|---|---|---|
| Use complement score | ✓ | When checked, complement score contributes to the calculation of the "raw score". |
| Use protease score | ✓ | When checked, protease score contributes to the calculation of the "raw score". |
| Use water loss score | ✓ | When checked, water-loss score contributes to the calculation of the "raw score". |
| Use ammonia loss score | ✓ | When checked, ammonia-loss score contributes to the calculation of the "raw score". |
| Use $A_2$ score | ✓ | When checked, $A_2$ score contributes to the calculation of the "raw score". |

#### Supplementary Table 2.

Description of the new de-novo associated columns in the “msmsScans.txt” output file from MaxQuant.

| Column name | Description |
|---|---|
| DN sequence | The de-novo AA regex sequence that corresponds to the one with the biggest raw score among all de-novo AA regex sequences for the specific MS/MS scan. |
| DN length | The length of the sequence that is stored in the column “DN sequence”. |
| DN min levenshtein distance | The minimum Levenshtein distance between the identified Andromeda sequence(column: “Sequence”) and the de-novo sequences resulting from the de-novo regex (column: “DN sequence”).<br>Levenshtein distance between two peptide sequences is the minimum number of single-amino acid edits (insertions, deletions, or substitutions) required to change one sequence into the other.<br>The MS/MS scans with no identified Andromeda sequence or no de-novo sequence or missing both will have DN min levenshtein distance equal to -1. |
| DN extension sequence | Sequence that was attached to the incomplete main path sequence in a second search for explaining the missing mass at a terminus. |
| DN extended | Whether or not a second search for explaining the missing mass at a terminus was successful. |
| DN complete | When marked with '+' the de-novo regex sequence (column: “DN sequence”) is full length with no mass missing.at the N- or/and C-terminal. |
| DN raw score | This is the score assigned to the de-novo regex sequence (column: “DN sequence”) and it is the sum of other 6 scores. Raw score = direct path score+complement score+protease score+a2 score+water-loss score+ammonia-loss score |
| DN extension score | Score that was achieved in a second search for explaining the missing mass at a terminus. |
| DN extension norm. score | Normalized score that was achieved in a second search for explaining the missing mass at a terminus. |
| DN (normalized) score | This is the raw score divided by the precursor mass. |

| Column name | Description |
|---|---|
| DN complete score | The complete score equals the normalized score, in case the sequence goes from terminus to terminus, i.e. is completely sequencing the peptide. Otherwise the complete score equals zero. |
| DN combined score | The combined score is a combination of the complete score and the gap score. Both of these scores are ranked, and then the sum of the two ranks is taken and normalized to lie between 0 and 100. |
| DN nterm mass | The mass that is missing at the N-terminal of the de-novo sequence tag. The sum of the de-novo sequence tag mass and the DN nterm mass is equal to the precursor mass associated with the specific MS/MS scan. |
| DN cterm mass | The mass that is missing at the C-terminal of the de-novo sequence tag. The sum of the de-novo sequence tag mass and the DN cterm mass is equal the precursor mass associated with the specific MS/MS scan. |
| DN missing mass | The sum of DN nterm mass and DN cterm mass. |
| DN nterm delta score | The score difference between the solution and the best solution that is not allowed to connect to the N-terminus. |
| DN cterm delta score | The score difference between the solution and the best solution that is not allowed to connect to the C-terminus. |
| DN term delta score | The maximum of the above two scores. |
| DN full length delta score | The difference in raw scores between the best and the second-best scoring solution. If there is no second-best solution, the gap score equals the raw score. |
| DN protease score | The reward for terminal parts of the sequence that are in an agreement with the specified protease that was used for generating peptides from proteins. |
| DN complement score | The score contribution of ions that are complementary to the ions in the direct path. |
| DN A2 score | The reward for the presence of an a2 ion. |
| DN isomer score | |
| DN water loss score | The reward for the presence of ions resulting from the loss of a water molecule in case the main path contains any of the amino acids D, E, S, or T. |
| DN ammonia loss score | The reward for the presence of ions resulting from the loss of an ammonia molecule in case the main path contains any of the amino acids K, N, Q, or R. |

| Column name | Description |
|---|---|
| DN agrees with andromeda | When marked with '+' the de-novo regex sequence (column: “DN sequence”) matches the sequence from the database search (column: “Sequence”) that passed the PSM FDR control (Column: “Identified”==”+”). Here a “match” is considered even if the de-novo regex sequence is a subsequence of the database search one. |
| DN agrees with andromeda complete | When marked with '+' the de-novo regex sequence (column: “DN sequence”) matches the sequence from the database search (column: “Sequence”) that passed the PSM FDR control (Column: “Identified”==”+”). Here a “match” is considered only the full-length match. |
| DN all sequences | All the de-novo AA regex sequences associated with a specific MS/MS scan separated by “;” sorted by their normalized score in descending order. |
| DN all scores | The normalized score of all the de-novo AA regex sequences associated with a specific MS/MS scan separated by “;” in descending order. |
| DN all agrees | “+” or “-” separated by “;” for each one of the de-novo regex sequences (column: “DN all sequences”) in the same order. When marked with '+' the de-novo regex sequence matches the sequence from the database search (column: “Sequence”). Here a “match” is considered even if the de-novo regex sequence is a subsequence of the database search one. |
| DN any agrees | When marked with '+' any of the de-novo regex sequences (column: “DN all sequences”) matches the sequence from the database search (column: “Sequence”). Here a “match” is considered even if the de-novo regex sequence is a subsequence of the database search one. |
| DN number of steps | The number of different steps that correspond to the path associated with the de-novo regex sequence (column: “DN sequence”). |
| DN is dominantly y | It is true if the main path corresponds to the y ion series. This is known for sequences that connect to at least one terminus. |

Three different kinds of brackets (round, square, and curly) can be found in a regex de-novo-sequence. The round brackets “()” contain the modification of the amino acid that is located on the left side of the brackets. For example M(Oxidation (M)). The square brackets “[]” contain two amino acids that their order cannot be distinguished by MaxNovo due to the absence of fragment ion peaks in the MS/MS scan. For example [WR] can be either WR or RW. The curly brackets “{}” contain amino acids that are separated by the pipe character “|”. The separated by “|” amino acids are equally possible to be at that position because they are isobaric or almost isobaric (up to mass tolerance error). For example {I|L} can be either I or L.
